# Adjustment for index event bias in genome-wide association studies of subsequent events

**DOI:** 10.1101/436063

**Authors:** Frank Dudbridge, Richard J. Allen, Nuala A. Sheehan, A. Floriaan Schmidt, James C. Lee, R. Gisli Jenkins, Louise V. Wain, Aroon D. Hingorani, Riyaz S. Patel

## Abstract

Following numerous genome-wide association studies of disease susceptibility, there is increasing interest in genetic associations with prognosis, survival or other subsequent events. Such associations are vulnerable to index event bias, by which selection of subjects according to disease status creates biased associations if common causes of incidence and prognosis are not accounted for. We propose an adjustment for index event bias using the residuals from the regression of genetic effects on prognosis on genetic effects on incidence. Our approach eliminates this bias when direct genetic effects on incidence and prognosis are independent, and otherwise reduces bias in realistic situations. In a study of idiopathic pulmonary fibrosis, we reverse a paradoxical association of the strong susceptibility gene *MUC5B* with increased survival, suggesting instead a significant association with decreased survival. In re-analysis of a study of Crohn’s disease prognosis, four regions remain associated at genome-wide significance but with increased standard errors.

## Introduction

The majority of genome-wide association studies (GWAS) conducted to date have studied susceptibility to disease. They have provided insights into biological mechanisms leading to disease, enabled Mendelian randomisation studies of risk factors, and shown promise for population screening ^1^. However, such studies are not necessarily informative on the course of disease, so their results cannot immediately be utilised to identify therapeutic targets or inform clinical management ^2^. Among a few published GWAS of survival, associated single nucleotide polymorphisms (SNPs) have tended to differ from those associated with susceptibility ^3-8^. With many collections of disease cases now genotyped by studies of susceptibility, more GWAS of severity, prognosis and survival are expected in the coming years.

Association studies of such subsequent events are vulnerable to index event bias, whereby biased associations can result from selection of subjects according to their disease status ^9^. This is one of several types of selection bias whose relevance to genetic epidemiology has recently been discussed ^10,11^. Independent causes of disease become correlated when selecting only the cases of disease, creating indirect associations between causes of disease with subsequent events (figure 1). A well-known example is the so-called obesity paradox whereby, among individuals with cardiovascular disease (CVD), those with higher body-mass index (BMI) tend to survive longer ^12^. A possible explanation is that, if an individual with CVD has a high BMI, they may well have lower levels of other risk factors. If those lower levels tend to increase survival, then increased BMI may be associated with longer survival. In the notation of figure 1, BMI plays the role of the SNP *G* while *X* is CVD and *Y* is survival. It remains controversial whether this paradox is explained by index event bias ^13^.

**Figure 1.**
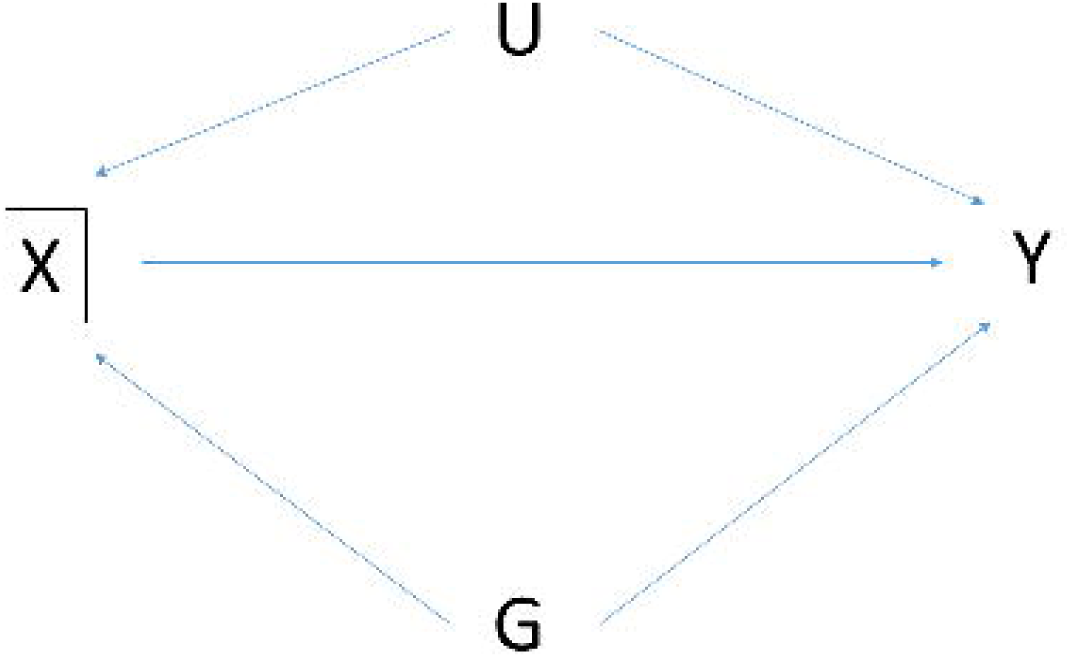
Directed acyclic graph of association of SNP G with prognosis *Y* conditional on incidence *X*. *U* is a composite variable including all common causes of *X* and *Y*, and may include polygenic effects as well as non-genetic risk factors. In our examples, *X* is idiopathic pulmonary fibrosis or Crohn’s disease, and *Y* is survival or prognosis. Conditioning on *X* induces the moralised association between *G* with *U* shown by the dotted line. This creates association between *G* and *Y* via the path *G* - *U* → *Y* in addition to the direct effect *G* → *Y*.

We will for simplicity refer to the index event as incidence, although our arguments also apply to selection or adjustment for a quantitative trait ^11^. Similarly we will refer to subsequent events as prognosis, with the understanding that this could refer to any phenotype subsequent to, and not a cause of, the index event.

In epidemiological studies, known confounders of incidence and prognosis have been used to construct propensity scores that effectively mitigate index event bias ^14^. Such approaches are difficult in genetic studies, because there may be a substantial polygenic confounder that cannot be modelled directly nor easily captured by a propensity score ^11^. Recently, the implications of index event bias have been discussed in the contexts of genetic association discovery ^15^ and Mendelian randomisation ^2^. Although the magnitude of bias appears small in currently typical settings, it is unclear how GWAS will be affected as studies increase in magnitude and polygenic analyses combine effects over thousands of variants.

Previously, expressions for index event bias have been derived when selecting on a binary disease trait ^13,16,17^ and when adjusting for a heritable covariate ^11^. These studies, however, have not identified methods for correcting this bias.

Some authors have considered bias when analysing a risk factor for the trait under selection ^17-20^. For example, accepting BMI as a cause of type-2 diabetes, a SNP with a direct effect on type-2 diabetes may have a biased association with BMI when studied within a case/control sample of type-2 diabetes (figure 2). This is different from the situation considered here as the trait of interest is a precursor of, and not subsequent to, the index event. This has also been called an index event bias ^17^, but here we reserve the term for when the trait of interest is subsequent to the selection criterion (figure 1). Methods are available to adjust analyses of risk factors for selection into case/control studies ^18,19,21,22^, but they do not apply here, when selection bias acts entirely through unobserved confounders.

**Figure 2.**
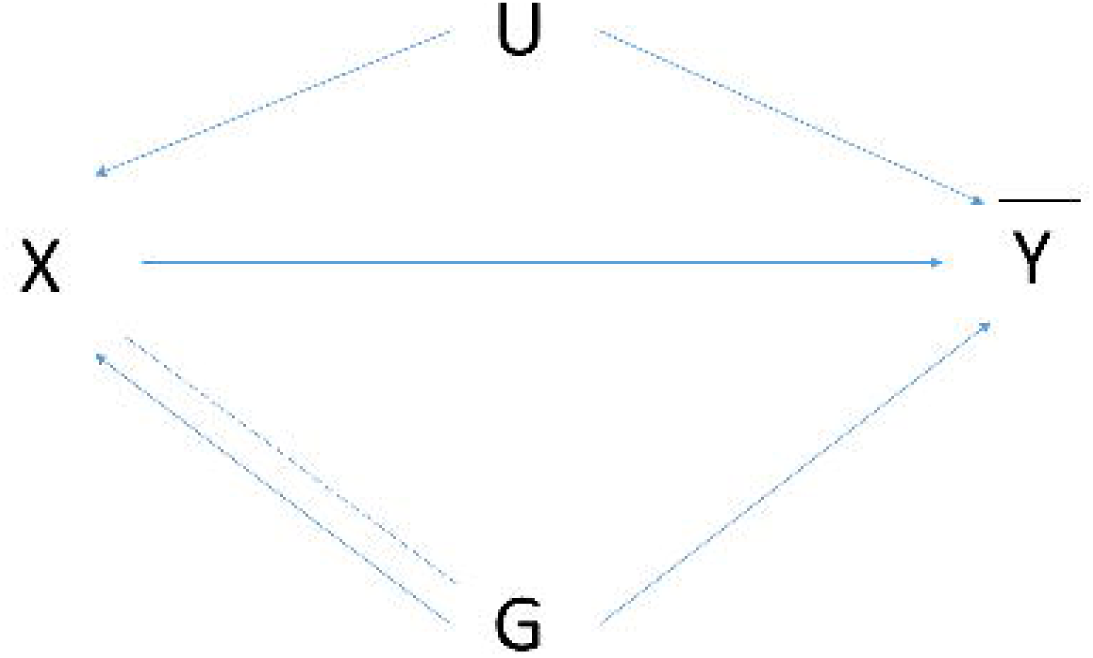
Directed acyclic graph of association of SNP *G* with risk factor *X* conditional on outcome *Y*. *U* is as in figure 1. For example, *X* may be body mass index, and *Y* may be type-2 diabetes, with the study design being case/control or a cohort depleted for cases, such as UK Biobank ^17^. Conditioning on *Y* induces the moralised association between *G* and *U* shown by the dotted line. This creates association of *G* with *X* via the path *G* - *U* → *X*, in addition to the direct effect *G* → *X*. The direct effect itself is biased by conditioning on *Y*, as shown by the additional dotted line connecting *G* and *X*. The resulting selection bias is not the focus of this paper.

Here we propose an adjustment for index event bias in GWAS of subsequent events. The main insight is that confounder effects are approximately constant across SNPs and can be estimated by regressing SNP effects on prognosis on SNP effects on incidence. We illustrate our approach in a GWAS of survival with idiopathic pulmonary fibrosis (IPF) and re-analyse a GWAS of Crohn’s disease prognosis.

## Results

### Adjustment for index event bias

For a single SNP, we assume that incidence *X* is linear in the coded genotype *G*, the combined common causes *U* of incidence and prognosis, and causes *E*_*X*_ unique to *X*:

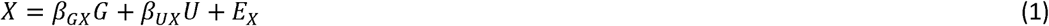

Similarly, assume that prognosis *Y* is linear in *G* and *U* with an additional main effect of *X*:

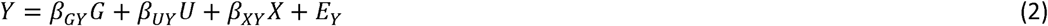

If *X* or *Y* are binary, we continue to argue from linear models by observing that logistic and probit link functions are approximately linear for small effects (Methods).

The effect of interest is the direct SNP effect on prognosis *β*_*GY*_, conditional on incidence *X* and confounders *U*. In practice however, the relevant confounders may not be observed and we can only estimate the SNP effect conditional on incidence, denoted by 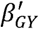. In the Methods we show that this estimand is the direct effect *β*_*GY*_ plus a bias that is linear in the effect on incidence *β*_*GX*_:

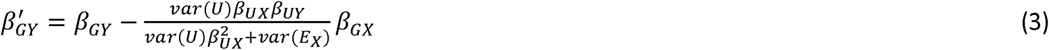

Notably, the coefficient of *β*_*GX*_ is negative if the confounder effects on incidence and prognosis, *β*_*UX*_ and *β*_*UY*_, have the same sign and positive if they have opposing signs.

Now consider Equation (3) applied to each one of a genome-wide set of SNPs. Assuming it has no interaction with each SNP, the non-genetic component of *U* is constant (figure 3). The genetic component of *U* equals the entire shared genetic basis of incidence and prognosis, minus any component due to the SNP under consideration. For polygenic traits, the variation explained by individual SNPs is small in relation to the total genetic variance, so we may assume that the genetic component of *U* is approximately constant across SNPs. Therefore we assume that 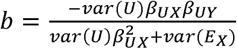 is approximately constant across SNPs, and may be obtained from the linear regression of 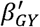 on *β*_*GX*_, giving the bias-corrected effects

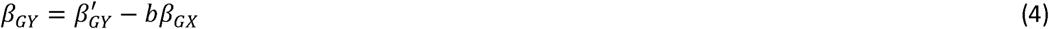

**Figure 3.**
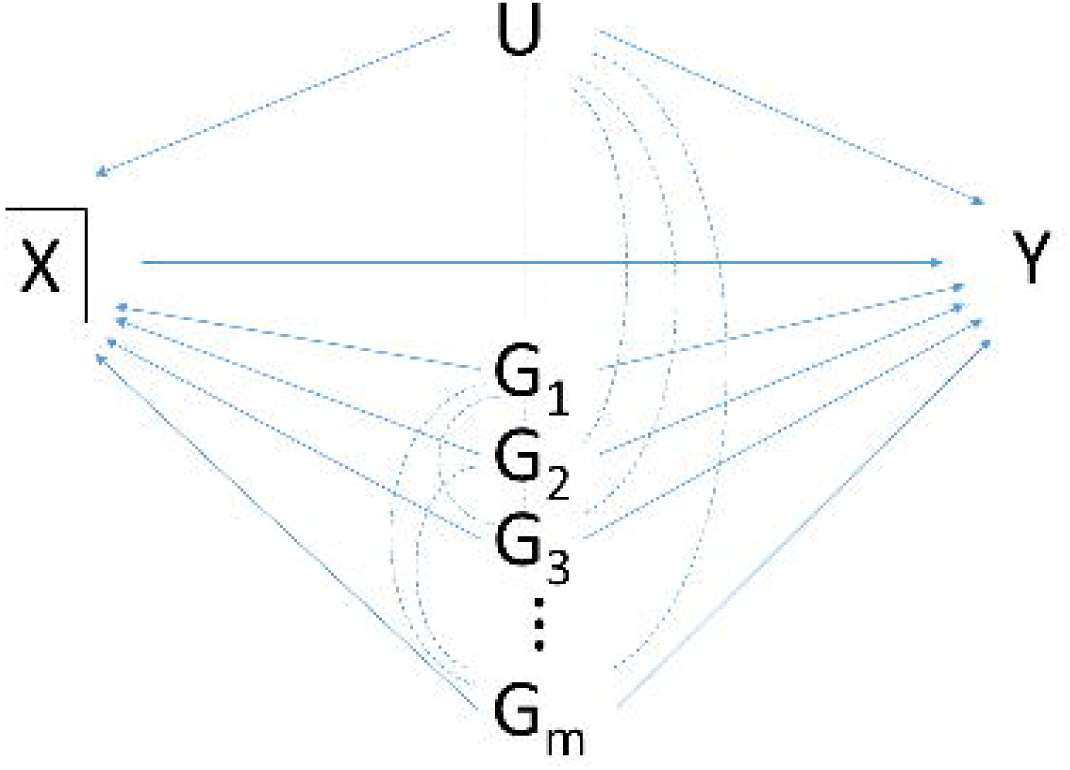
Directed acyclic graph of association of SNPs *G*_*i*_ with prognosis *Y* conditional on incidence *X*. *U* is as in figure 1. Conditioning on *X* induces the moralised associations shown by dotted lines. These create association of each *G*_*i*_ with *Y* via the path *G*_*i*_ -*U* → *Y* and all paths *G*_*i*_ - *G*_*f*_ → *Y* where *i* ≠ *j*. Under a polygenic model in which individual SNPs explain little covariation between *X* and *Y*, the combined effect of *U* and all *G*_*j*≠*i*_ is approximately constant across SNPs *G*_*i*_. If a SNP *G*_*k*_ has a major effect on *X* and/or *Y*, the associations of *G*_*j*≠*k*_ can be conditioned on *G*_*k*_ to prevent the major gene contributing to index event bias.

In practice we have finite sample estimates 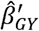 and 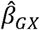 and the regression will yield an estimate 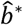 that is biased towards zero by sampling error in 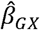, consequently under-correcting in equation (4). In the Methods we describe two approaches to adjust for this regression dilution. The first obtains a bias-reduced estimate of *b* as 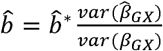, where *var*(*β*_*GX*_) is approximated by the Hedges-Olkin estimator 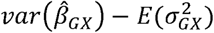 with 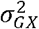 the squared standard error of 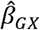. This simple adjustment is sufficiently accurate for simulation studies. For the analysis of data, we developed an improved version of the simulation extrapolation (SIMEX) algorithm ^23,24^. This is more computationally intensive but also more accurate, and yields confidence intervals for *b* so that 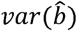 can be estimated. Details are given in the Methods and Supplementary Note 1.

From Equation (4), the variance of the bias-adjusted estimate is approximately

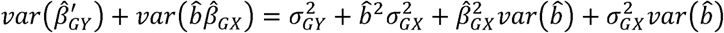

Although there is no theory that 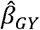 is normally distributed, a normal approximation works well in practice. Further details are provided in the Methods.

Summarising, we propose the following procedure to correct index event bias in GWAS of prognosis:

1. For each SNP, obtain its estimated effect on incidence 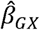 with standard error *σ*_*GX*_, and its estimated effect on prognosis 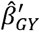 conditional on *x* with standard error *σ*_*GY*_.
2. Obtain 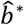 as the slope of the linear regression of 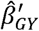 on 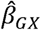.
3. Adjust 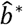 for regression dilution using SIMEX (in data) or Hedges-Olkin adjustment (in simulations), obtaining the corrected slope estimate 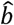.
4. For each SNP, the bias-corrected estimate of its effect on prognosis is 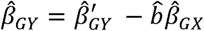 with standard error 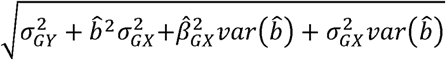.
5. Hypothesis tests and *p*-values for each SNP may be computed by referring the ratio of the adjusted estimate and its standard error to the standard normal distribution.

In the Methods we discuss some assumptions upon which this procedure is based. One implication is that the regression of step 2 should be performed on independent SNPs. Therefore we prune GWAS SNPs by linkage disequilibrium (LD) as a pre-processing step. However, this is only required to obtain a valid estimate of the slope *b*, which once obtained can be applied to all SNPs.

If there are major genes accounting for substantial covariation in *x* and *y*, the genetic component of *u* may not be constant across SNPs and the assumption of a constant regression slope *b* is violated. This problem could be avoided by conditioning the prognosis associations on the major genes, thereby estimating the bias through all confounders except those genes. The resulting correction is then appropriate for all SNPs including those in major genes (figure 3). A similar approach can be taken to polygenic scores aggregating the small effects of several individual SNPs.

Our most serious assumption is no correlation between effects on incidence *β*_*GX*_ and direct effects on prognosis *β*_*GY*_, for those SNPs entering the regression of step 2. If incidence and prognosis have common biological mechanisms then this assumption may be violated and create bias in *b* and hence in 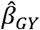. However, considering pleiotropy in general some authors have argued that independence of effects is likely to be the norm in complex disease ^25^. We explore this assumption in the following simulations and return to this point in the Discussion.

### Simulations

Firstly we simulated 100,000 independent SNPs of which 5000 (5%) had effects on incidence only, 5000 had effects on prognosis only and 5000 had effects on both incidence and prognosis. Incidence and prognosis were simulated as quantitative traits under additive models with 50% heritability (Methods), with a non-genetic confounder (representing the combined effects of all such factors) simulated to explain 40% of variation in both incidence and prognosis. No direct effect of incidence on prognosis was simulated (*β*_*XY*_ = *0*). Data were simulated for 20,000 unrelated individuals. Incidence and prognosis were analysed as quantitative traits using linear regression, with the prognosis model adjusting for incidence as a covariate. This simulation, which reflects the scenario discussed by Aschard et al ^11^, satisfies the assumptions of our procedure while creating a high degree of index event bias (Methods).

Table 1 shows type-1 error rates for the standard unadjusted analysis and for the adjusted analysis using our procedure. For all analyses the rate is close to the nominal level when averaged over all SNPs; however, the majority of SNPs have no index event bias. Among the SNPs with effects on incidence, the type-1 error is inflated for the unadjusted analysis while our approach achieves the correct rate when there is no genetic correlation between incidence and prognosis. Under positive genetic correlation, the type-1 error increases for all analyses, but is consistently lower for our adjusted analysis. For some individual SNPs the type-1 error can be very high under the unadjusted analysis but is substantially reduced under our approach, and at the nominal level when there is no genetic correlation. Under strong negative genetic correlation, our approach has slightly increased type-1 error compared to the unadjusted analysis. However this situation, if not implausible, is arguably less likely than positive genetic correlation ^2^. Results for the family-wise error were more pronounced and followed the same pattern.

**Table 1.** Power for quantitative incidence and prognosis with non-genetic confounding. Estimates shown as % with *P* < 0.05 over 1000 simulations of 100,000 independent SNPs. 5000 SNPs have effects on incidence only, 5000 on prognosis only and 5000 on both incidence and prognosis. Heritability of both incidence and prognosis is 50% with the genetic correlation shown over all SNPs. Common non-genetic factors explain 40% of variation in both incidence and prognosis. Rows 3-6 show type-1 error rates. All SNPs, mean power across the relevant SNPs. Family-wise error, probability of at least one SNP with effect on incidence but not on prognosis having 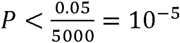. SNP with greatest increase (decrease) in power compares the adjusted analysis to the unadjusted.

|  |  |  |  |  |  |  |  |  |  |  |
| --- | --- | --- | --- | --- | --- | --- | --- | --- | --- | --- |
| Genetic correlation | 0 | 0 | .25 | .25 | .45 | .45 | -.25 | -.25 | -.45 | -.45 |
| Adjustment | No | Yes | No | Yes | No | Yes | No | Yes | No | Yes |
| All SNPs not affecting prognosis | 5.12 | 5.00 | 5.25 | 5.06 | 5.42 | 5.23 | 5.05 | 5.04 | 5.02 | 5.10 |
| All SNPs affecting incidence but not prognosis | 7.24 | 5.03 | 9.59 | 6.06 | 12.5 | 9.15 | 5.93 | 5.65 | 5.38 | 6.85 |
| SNP with highest type-1 error | 33.0 | 5.7 | 61.7 | 19.1 | 87.9 | 63.7 | 20.0 | 15.0 | 10.8 | 28.5 |
| Family-wise type-1 error | 22.3 | 5.5 | 61.0 | 12.8 | 94.8 | 53.4 | 12.1 | 10.0 | 6.8 | 16.3 |
| All SNPs affecting prognosis | 19.5 | 16.7 | 18.7 | 18.0 | 16.6 | 17.2 | 19.3 | 13.8 | 18.7 | 10.9 |
| All SNPs affecting incidence and prognosis | 20.3 | 16.5 | 16.7 | 16.5 | 10.0 | 11.9 | 21.3 | 13.1 | 20.6 | 8.38 |
| SNP with greatest increase in power | 6.6 | 39.2 | 18.0 | 50.1 | 34.8 | 59.5 | 6.9 | 34.8 | 5.2 | 14.9 |
| SNP with greatest decrease in power | 72.3 | 19.9 | 75.0 | 41.9 | 20.4 | 12.1 | 93.6 | 19.2 | 96.1 | 22.0 |

Table 1 also shows power for the same simulations. There is overall a modest drop in power for our approach, except under strong positive genetic correlation where there is a small increase. The power loss is greatest under strong negative genetic correlation. For individual SNPs, substantial differences can occur between methods. The most extreme cases entail a greater gain in power for the unadjusted analysis than for our approach, although this must be offset against the increased type-1 error. Supplementary tables 1 and 2 show absolute bias and mean square error. The pattern is similar in that the adjusted analysis has less bias, although this is offset by its increased standard error so that the differences in mean square error are barely discernible.

**Table 2.** Power for quantitative incidence and prognosis without non-genetic confounding. Parameters are as in table 1 except that there are no common non-genetic factors of incidence and prognosis.

|  |  |  |  |  |  |  |  |  |  |  |
| --- | --- | --- | --- | --- | --- | --- | --- | --- | --- | --- |
| Genetic correlation | 0 | 0 | .25 | .25 | .45 | .45 | -.25 | -.25 | -.45 | -.45 |
| Adjustment | No | Yes | No | Yes | No | Yes | No | Yes | No | Yes |
| All SNPs not affecting prognosis | 5.00 | 5.00 | 5.01 | 5.04 | 5.03 | 5.13 | 5.01 | 5.03 | 5.03 | 5.12 |
| All SNPs affecting incidence but not prognosis | 5.00 | 5.01 | 5.17 | 5.72 | 5.61 | 7.30 | 5.18 | 5.64 | 5.61 | 7.20 |
| SNP with highest type-1 error | 8.02 | 8.02 | 8.50 | 15.2 | 13.0 | 35.6 | 8.30 | 13.7 | 13.4 | 33.7 |
| Family-wise type-1 error | 4.50 | 3.90 | 4.90 | 8.80 | 8.00 | 24.7 | 5.30 | 9.20 | 7.80 | 22.1 |
| All SNPs affecting prognosis | 16.6 | 16.6 | 16.2 | 15.6 | 15.1 | 13.3 | 16.2 | 15.6 | 15.1 | 13.3 |
| All SNPs affecting incidence and prognosis | 16.4 | 16.4 | 15.4 | 14.4 | 13.0 | 9.75 | 15.4 | 14.5 | 12.9 | 9.86 |
| SNP with greatest increase in power | 31.2 | 33.3 | 25.1 | 36.4 | 9.20 | 13.9 | 18.6 | 29.4 | 6.70 | 10.7 |
| SNP with greatest decrease in power | 20.1 | 17.9 | 53.9 | 35.4 | 62.0 | 26.2 | 46.1 | 31.9 | 56.6 | 26.2 |

We then repeated the simulation with no non-genetic confounding, so that bias only arises through genetic correlation violating our independence assumption. Table 2 shows that type-1 error for our approach is similar to that when non-genetic confounding is present, but for the unadjusted analysis the errors are reduced and generally closer to the nominal level than for our approach. Again there is a slight decrease in power under our approach, with considerable increases and decreases possible for individual SNPs. Supplementary tables 3 and 4 show similar patterns for absolute bias and mean square error.

Tables 3 and 4 and supplementary tables 5 to 8 show similar patterns when the incidence and prognosis traits are binary and prognosis is analysed in cases only (Methods). Supplementary tables 9 to 12 also show similar patterns when the prognosis is quantitative and analysed either in cases only or in the full sample with adjustment for case/control status (Methods). These results confirm that our approach is applicable when incidence is analysed by logistic regression, and that it maintains the correct type-1 error rate when there is no genetic correlation between incidence and prognosis, and otherwise has a lower type-1 error rate than the unadjusted analysis except under strong negative genetic correlation or no non-genetic correlation. While the relative strength of genetic and non-genetic confounding is unknown in practice, we might expect them to act in the same direction, and the genetic confounding not to dominate the non-genetic. These are the scenarios in which our approach does best; furthermore the type-1 errors are more consistent under different scenarios under our approach than the unadjusted analysis. Turning to power, there is again a modest reduction in general, with more substantial gains and losses possible for individual SNPs. Overall we conclude that our approach can be preferred to an unadjusted analysis.

**Table 3.** Power for binary incidence and prognosis with non-genetic confounding. Parameters are as in table 1 with cases defined as subjects in the top 20^th^ percentile of the incidence trait, and poor prognosis as cases in the top 50^th^ percentile of the prognosis trait. Prognosis is analysed in cases only.

|  |  |  |  |  |  |  |  |  |  |  |
| --- | --- | --- | --- | --- | --- | --- | --- | --- | --- | --- |
| Genetic correlation | 0 | 0 | .25 | .25 | .45 | .45 | -.25 | -.25 | -.45 | -.45 |
| Adjustment | No | Yes | No | Yes | No | Yes | No | Yes | No | Yes |
| All SNPs not affecting prognosis | 5.02 | 5.00 | 5.04 | 5.01 | 5.05 | 5.03 | 5.01 | 5.01 | 5.00 | 5.03 |
| All SNPs affecting incidence but not prognosis | 5.37 | 5.02 | 5.66 | 5.21 | 5.96 | 5.68 | 5.18 | 5.21 | 5.05 | 5.60 |
| SNP with highest type-1 error | 11.6 | 8.20 | 13.7 | 8.5 | 18.2 | 14.6 | 8.40 | 9.20 | 8.30 | 14.0 |
| Family-wise type-1 error | 5.80 | 5.60 | 7.70 | 5.20 | 12.0 | 8.80 | 4.60 | 6.20 | 6.60 | 8.70 |
| All SNPs affecting prognosis | 8.99 | 8.63 | 8.32 | 8.39 | 7.49 | 6.48 | 9.28 | 8.16 | 9.29 | 7.34 |
| All SNPs affecting incidence and prognosis | 9.13 | 8.59 | 7.80 | 8.00 | 10.9 | 9.60 | 9.74 | 7.81 | 9.77 | 6.33 |
| SNP with greatest increase in power | 36.5 | 48.1 | 24.5 | 34.6 | 23.0 | 29.9 | 7.20 | 16.6 | 5.90 | 8.50 |
| SNP with greatest decrease in power | 51.4 | 33.3 | 33.0 | 27.0 | 10.9 | 9.60 | 50.4 | 20.7 | 63.2 | 20.9 |

**Table 4.** Power for binary incidence and prognosis without non-genetic confounding. Parameters as in table 3 except that that there are no common non-genetic factors of incidence and prognosis.

|  |  |  |  |  |  |  |  |  |  |  |
| --- | --- | --- | --- | --- | --- | --- | --- | --- | --- | --- |
| Genetic correlation | 0 | 0 | .25 | .25 | .45 | .45 | -.25 | -.25 | -.45 | -.45 |
| Adjustment | No | Yes | No | Yes | No | Yes | No | Yes | No | Yes |
| All SNPs not affecting prognosis | 5.00 | 5.00 | 5.00 | 5.01 | 5.00 | 5.03 | 5.00 | 5.01 | 5.00 | 5.03 |
| All SNPs affecting incidence but not prognosis | 5.00 | 5.01 | 5.02 | 5.22 | 5.10 | 5.70 | 5.03 | 5.21 | 5.11 | 5.66 |
| SNP with highest type-1 error | 7.30 | 7.30 | 7.70 | 8.60 | 7.70 | 15.5 | 8.30 | 9.10 | 8.10 | 14.4 |
| Family-wise type-1 error | 3.50 | 3.70 | 4.40 | 5.20 | 5.60 | 10.1 | 4.80 | 5.50 | 5.60 | 9.10 |
| All SNPs affecting prognosis | 8.76 | 8.77 | 8.60 | 8.38 | 8.25 | 7.58 | 8.61 | 8.41 | 8.25 | 7.63 |
| All SNPs affecting incidence and prognosis | 8.71 | 8.72 | 8.38 | 8.00 | 7.66 | 6.45 | 8.39 | 8.04 | 7.66 | 6.51 |
| SNP with greatest increase in power | 13.6 | 14.9 | 13.9 | 18.1 | 5.90 | 7.80 | 8.40 | 12.5 | 5.80 | 7.80 |
| SNP with greatest decrease in power | 44.2 | 42.5 | 53.3 | 42.6 | 48.9 | 27.7 | 51.9 | 43.4 | 43.1 | 25.8 |

### Idiopathic pulmonary fibrosis

A recent GWAS meta-analysis of idiopathic pulmonary fibrosis (IPF) confirmed the strong association of mucin 5B (*MUC5B*) with incidence^26^. We re-analysed 612 UK cases and 3,366 UK controls that we had contributed to that meta-analysis, obtaining an odds ratio of 5.64 for the SNP rs35705950 in *MUC5B* (95%CI: 2.73, 6.72; Wald test *P*=2.9e-83), and conducted a GWAS of survival time in 565 of the cases (Methods). Similar to previous studies ^27^ the risk allele of rs35705950 was associated with increased survival in our study (hazard ratio 0.766; 95%CI: 0.634, 0.925; Wald test *P*=0.0057). This apparently paradoxical result could arise from index event bias, given the strong odds ratio for incidence. We applied our regression based adjustment using 140,092 LD-pruned SNPs with imputation *R*^2^ ≥ 0.99 (Methods). Here we focus only on the effect of rs35705950 in *MUC5B*; full results of the survival GWAS will be reported separately.

The regression of survival log hazard ratios on incidence log odds ratios gave a coefficient of −0.025. The sign of this coefficient changed under the Hedges-Olkin based adjustment for regression dilution, which is implausible because regression dilution bias is the ratio of two variances^28^. However applying our SIMEX based adjustment, the coefficient decreased to −65.63 (95% CI: −65.88, −5.68). The very wide confidence interval reflects high standard errors on individual SNP effects. Nevertheless the coefficient is significantly negative, which implies that there are common causes of incidence and prognosis that have the same net direction of effect.

The asymmetry in the confidence interval suggests that a normal approximation for 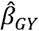 would be inappropriate. We therefore generated an empirical distribution of 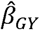 to assess its significance (Supplementary Note 1). Of 10,000 simulations of 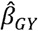, none were less than zero, suggesting that rs35705950 has a positive log hazard ratio with *P*-value of order less than 10^−4^. The empirical 95% confidence interval for the log hazard ratio was (11.58, 126.56), suggesting an extremely strong effect on survival. However, in view of the substantial uncertainty in 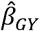 we refrain from drawing a strong conclusion beyond the direction of effect.

Given the strong effect of *MUC5B* on incidence, it is possible that it dominates the estimation of index event bias and that our approach over-or under-corrects the bias in *MUC5B* itself (figure 3). We therefore repeated the analysis after conditioning the survival SNP effects on rs35705950 genotype. The results were very similar, with the regression of survival effects on incidence effects now giving a coefficient of −0.028, decreasing to −59.79 (95%CI: −59.87, −10.52) after correcting for regression dilution.

We repeated our simulations using a sample size of 612 cases and 3366 controls, generating survival times from the exponential model using the simulated prognosis trait as the log hazard, and testing association using Cox regression (supplementary tables 13 and 14). Adjusted and unadjusted analyses had similar overall properties, suggesting that our approach could be applicable in this setting. Together our results suggest that the paradoxical association of *MUC5B* with increased survival could indeed be due to index event bias, and that the risk allele of *MUC5B* is in fact associated with decreased survival.

### Crohn’s disease

Odds ratios for Crohn’s disease have been published by the International Inflammatory Bowel Disease Genetics Consortium^29^ and for prognosis (binary good/poor) by a subsequent study by Lee et al^6^. The latter study identified four regions associated with prognosis at genome-wide significance (*P* < 5 × 10^−8^), none of which were significantly associated with disease susceptibility.

We reanalysed the summary statistics using our regression-based adjustment with 29,715 LD-pruned SNPs with imputation *R*^2^ ≥ 0.99 (Methods). The regression of prognosis log odds ratios on incidence log odds ratios gave a coefficient of −0.042, which decreased to −0.264 with SIMEX adjustment for regression dilution (95%CI: −0.299, −0.236). Here the Hedges-Olkin based adjustment gave a similar result of −0.272. Again the negative coefficient implies that there are common causes of incidence and prognosis with concordant directions of effect.

After adjusting the association of each SNP, the genomic control inflation parameter was 1.016 compared to 1.024 in the unadjusted analysis. Of the four reported associations with prognosis, three of the lead SNPs remained genome-wide significant, while association of the lead SNP in the *MHC* region was attenuated to just short of genome-wide significance. However, another *MHC* SNP, which was genome-wide significant in the index study, did remain so after adjustment (table 5). Following Lee et al^6^ we inspected the associations with prognosis of 170 SNPs robustly associated with incidence (supplementary data 1). None of these SNPs were significantly associated with prognosis, after correcting for 170 tests.

**Table 5.** *P*-values for four regions associated with Crohn’s disease prognosis. Unadjusted P, Wald test P-value reported by Lee et al ^6^. Adjusted *P*, Wald test *P*-value from our adjusted analysis. The lead variant in *MHC* in Lee et al, rs9279411 does not achieve genome-wide significance in our adjusted analysis but is approximately 72kb proximal to rs114575504, which does achieve significance.

| Chromosome | Mb | Variant | Nearest gene | Unadjusted *P* | Adjusted *P* |
| --- | --- | --- | --- | --- | --- |
| X | 112.9 | rs5929166 | *XACT* | 4.56e-9 | 6.56e-9 |
| 6 | 31.7 | rs9279411 | *MHC* | 5.46e-9 | 7.93e-8 |
| 6 | 31.7 | rs114575504 | *MHC* | 9.37e-9 | 4.46e-8 |
| 6 | 109.0 | rs3778586 | *FOXO3* | 1.47e-8 | 2.66e-8 |
| 7 | 45.9 | rs75764599 | *IGFBP1* | 4.32e-8 | 4.00e-8 |

Previously, Lee et al reported a negative genetic correlation between Crohn’s disease incidence and prognosis^6^. This is consistent with the negative coefficient in our regression of prognosis effects on incidence effects, and so could be explained by index event bias. However, our adjusted estimates of prognosis effects are by construction uncorrelated with the incidence effects, and so any analysis of genetic correlation based on our adjustments would be misleading.

## Discussion

Awareness of index event and related biases ^2,10,11^ has grown as attention turns to follow-up of GWAS. Our interest in this problem arose within the GENetIcs of sUbSequent Coronary Heart Disease (GENIUS-CHD) consortium ^30^, which aims to identify risk factors for recurrent coronary events in patients with coronary heart disease. In a simulation study ^15^, we showed that index event bias could be small in GWAS. Here we have confirmed this for Crohn’s disease, but have shown an example in IPF where a strong effect on susceptibility appears to create a substantial bias that reverses the survival effect. This illustrates how index event bias can have variable effects in different studies, and reinforces the need to adjust for it to be confident in any genetic associations with prognosis.

The critical assumption of our approach is that direct genetic effects on prognosis are independent of those on incidence. Since GWAS of susceptibility have been motivated by the discovery of novel treatment targets, our assumption may seem incompatible with the premise of GWAS. Indeed, shared pathways of incidence and prognosis have been observed in coronary heart disease, in which statins have proved effective in preventing both initial and recurrent events ^31^. For phenotypes related to cumulative effects of long term exposures, such as CVD but also perhaps some psychiatric traits, such shared pathways may be common. But for conditions in which prognosis depends upon the response to an initiating event, as perhaps in cancer or infectious diseases, the determinants of prognosis are conceivably independent of those for incidence. Even in CVD, determinants of arterial plaque development may be independent of those for plaque rupture. For immune-mediated disease, where a break in immunological tolerance is the key event at disease initiation, it is expected that other pathways drive disease course, since tolerance can only be broken once to a particular antigen. Also, where developmental mechanisms contribute to predisposition to late-onset disease, the determinants of prognosis may plausibly be independent. Some have argued that independently pleiotropic effects are likely to be typical for complex disease ^25^: for most pairs of traits the genetic effects on the first are independent of the corresponding effects on the other. However, our independence assumption precludes any meaningful analysis of genetic correlation between incidence and prognosis.

Our simulations showed reduced type-1 error rates for our procedure compared to an unadjusted analysis, except in the case of strong negative genetic correlation between incidence and prognosis or no non-genetic correlation. Power is slightly reduced overall but may be considerably increased for some individual SNPs. Again our approach performed more poorly under strong negative genetic correlation, which is arguably less likely than positive correlation. We simulated genetic architectures that were typical of complex diseases^32^ while allowing a high degree of index event bias. In smaller studies however, such as our IPF survival GWAS, our adjustment may have high variance resulting in more severely reduced power.

Our analysis of IPF suggests that a paradoxical association of the strong risk locus *MUC5B* with increased survival may be due to index event bias, and that in fact this gene may well cause decreased survival. It has been hypothesised that carriers of *MUC5B* risk alleles experience a milder form of disease, in line with the clinical heterogeneity of IPF^27^. While associations with prognosis can be explained by disease heterogeneity, they remain susceptible to index event bias whether or not disease subtype is accounted for (figure 4). It is therefore important to account for the bias to inform the interpretation of genetic associations with prognosis. Here, the reversal of direction for the *MUC5B* survival effect is largely due to its exceptionally high odds ratio for susceptibility. However our result, while significant, is imprecise and based on a sample that is small by current standards. It is crucial to replicate this result in larger samples or meta-analyses.

**Figure 4.**
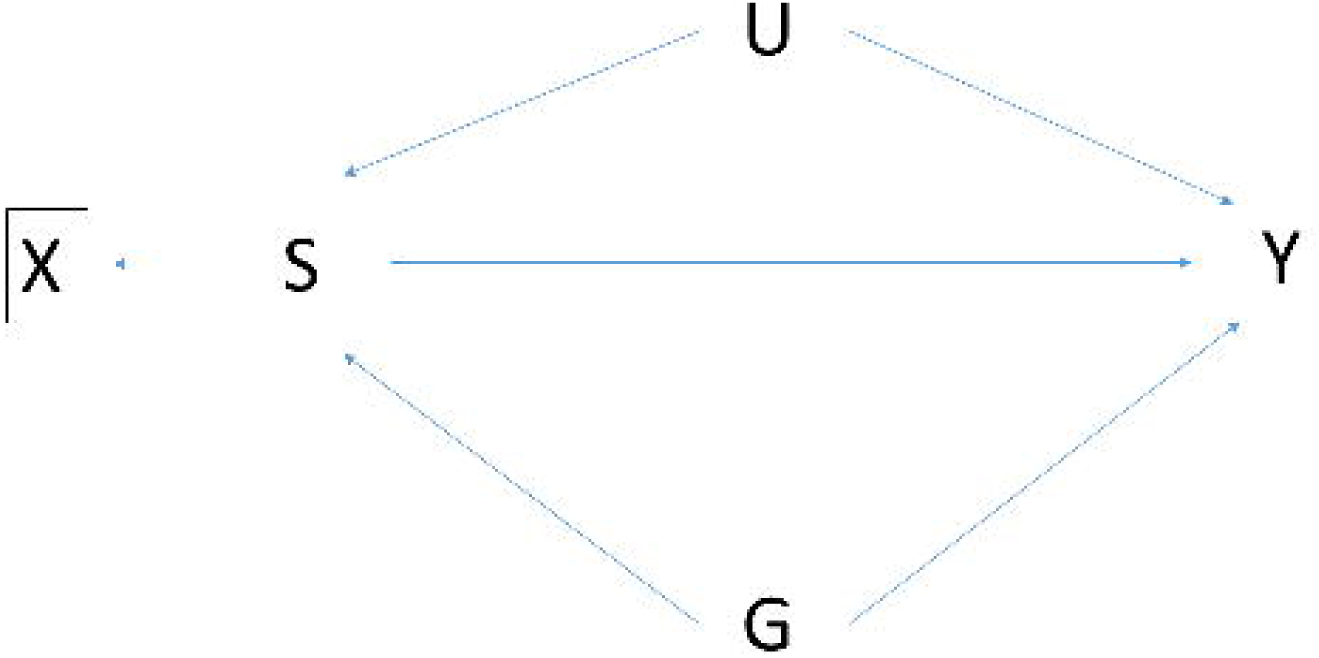
Association of SNP *G* with prognosis *Y* conditional on incidence *X* derived from trait *S*. *U* is a composite variable as in figure 1. For example, *X* may be a diagnosis of disease (eg Crohn’s disease), and *S* a subtype of disease (eg ileal, colonic, ileocolonic or healthy). Conditioning on *X*, which is a descendant of the collider *S*, induces the moralised association between *G* and *U* shown by the dotted line. This creates association of *G* with *Y* via the path *G* - *U* → *Y* in addition to its direct effect via *G* → *Y* and its mediated effect via *G* → *S* → *Y*. Further conditioning on S blocks the mediation path *G* → *S* → *Y* but leaves open the path *G* - *U* → *Y* creating index event bias.

We confirmed the genome-wide significance of four regions previously associated with Crohn’s disease prognosis, but did not identify any further associations with prognosis. In their index study, Lee et al^6^ adjusted for disease location before inspecting associations between disease prognosis and 170 susceptibility SNPs. This was done because some of the criteria used to define severe disease (e.g. need for recurrent surgery) could lead to an over-representation of patients with ileal disease, for whom surgery is more commonly used because the operation carries lower morbidity than colonic surgery and does not leave a permanent stoma. Considering location as a disease subtype, adjustment for location might modify the prognosis associations for SNPs with effects on particular disease locations (figure 4). We did not adjust for location here, as our aims were to identify associations with prognosis independently of possible mechanism, and to demonstrate the utility of our approach on published summary statistics.

Our procedure has some similarity to Egger regression applied to Mendelian randomisation (MR-Egger)^33^. Both procedures assume the structure in figure 1, regress SNP effects on one trait on SNP effects on another, and require an independence assumption. However, while the focus of MR-Egger is on the slope of the regression (the causal effect of exposure on outcome), and on the intercept (the magnitude of directional pleiotropy), our focus here is on the residuals, which provide the adjusted effect estimates when added to the intercept.

We may draw on the analogy with MR-Egger to contemplate other approaches based on the ratio of prognosis effects to incidence effects. Such approaches would entail other assumptions that require careful consideration. For example, a counterpart of the median ratio estimator ^34^ would assume that at least half of the SNPs considered have no direct effect on prognosis. Alternative approaches related to parallel work in Mendelian randomisation are a promising area for further development.

Our approach is robust to the use of the same subjects in the prognosis GWAS as in the incidence GWAS. This is because any correlation in prognosis and incidence phenotypes is by definition included in *U* and is therefore accounted for by our regression procedure. Indeed our simulations used the same subjects in both GWAS, and obtained the correct type-1 error rates when expected.

Our analytic result is derived from linear regression models, and is inexact for traits generated under other models ^13,17^. However, in practice our approach only requires that the bias is linear in the incidence effect, which we argue is approximately true for polygenic traits. This linear relationship is estimated from data, and while our theory provides an interpretation for it under some assumptions, our approach requires only that such a relationship exists. The data in our examples used log odds ratios and log hazard ratios, and our simulations suggested the linear approximation was acceptable in these cases.

When the incidence trait is binary, we have mainly considered a case-only analysis of prognosis. Other approaches are possible, such as setting the prognosis to a degenerate value for controls and then analysing cases and controls together, with adjustment for case/control status ^18^. In our simulations we found no systematic difference between the case/control and case-only analysis. Note that our approach could be applied in conjunction with the case/control analysis, and possibly with further adjustment for measured confounders of incidence and prognosis. This would have the desirable effect of reducing index event bias through several complementary approaches at once.

We have not considered the case in which the trait of interest is a precursor of the trait under selection (figure 2). Selection bias also occurs in this case ^10,17,20^ but cannot immediately be corrected by our approach because it would require knowledge of the effects of all confounders (Supplementary Note 2). Methods exist to adjust for selection in this situation ^18,19,21,22^, although they do not allow for unmeasured confounders. It may be possible to combine our approach with these methods to more fully account for selection bias in this situation.

Other forms of selection bias may be present that are not addressed by our approach. For example, participation in either incidence or prognosis GWAS is often conditional on survival until time of recruitment, but there may be unmeasured common determinants of survival and incidence/prognosis that create further biases. We have previously shown survival bias to be potentially of similar importance to index event bias ^15^, and this should be borne in mind when performing studies of prognosis, particularly when the index event may be acute as in coronary heart disease. Censoring after diagnosis, for example from death by competing risks, may also create bias if there are common determinants of incidence, censoring and/or prognosis. Our approach is developed under a simple model of incidence and prognosis, but provides a starting point for extensions that model the disease course more precisely.

Our approach could be applied to correct index event bias of non-genetic exposures. If the effects of all polygenic SNPs are estimated conditional on the non-genetic exposure, we can estimate the bias through all confounders other than that exposure. The effect of the exposure can then be adjusted in the same way as for the SNPs, giving a new and potentially wide application for GWAS data.

We have focussed on reducing bias in estimating the direct effects of SNPs on prognosis, to gain insight into mechanisms of prognosis. A different goal may be to build prediction models of prognosis. In that case it is preferable to work with the unadjusted effects since they do represent the total associations with prognosis conditional on incidence.

We have proposed an approach to adjust for index event bias in GWAS of subsequent events that achieves unbiased results under an independence assumption and otherwise compares favourably with the unadjusted analysis. It integrates the identification and adjustment of the bias in a single statistical procedure. We believe this method can be recommended as a standard analysis for GWAS of subsequent events.

## Methods

### Bias adjustment

Recall that we assume incidence *X* is linear in the coded genotype *G*, the combined common causes *U* of incidence and prognosis, and causes *E*_*X*_ unique to *X* (equation 1):

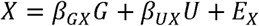

Similarly, assume that prognosis *Y* is linear in *G, U* and *X* (equation 2):

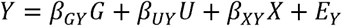

These are not necessarily causal models, but reflect a parameterisation of associations between *G, U, X* and *Y* that is natural when the conditional independence structure is as in figure 1 without conditioning on *X*. We assume that *G, U, E*_X_ and *E*_y_ are pairwise uncorrelated and have no interactions in the models for *X* and *Y*. Polygenic effects may contribute to *U, E*_*X*_ and *E*_*Y*_.

Assume without loss of generality that *G, U, E*_X_ and *E*_*Y*_ each have mean zero and hence also *E*(*X*) = *E*(*Y*) = 0. Let 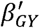 be the effect of *G* on *Y* conditional on *X*, but not on *U*. If 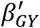 is estimated from the linear regression model

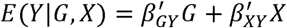

then the asymptotic ordinary least squares estimate is

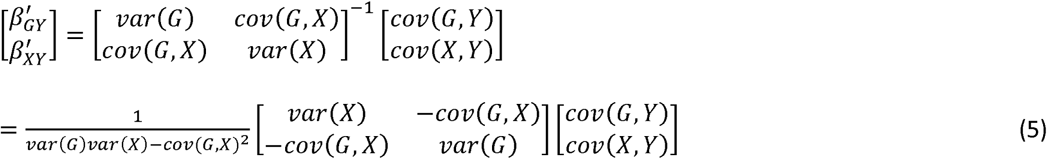

From Equation (1), under the assumptions above,

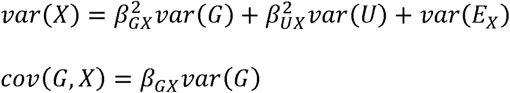

From Equation (2),

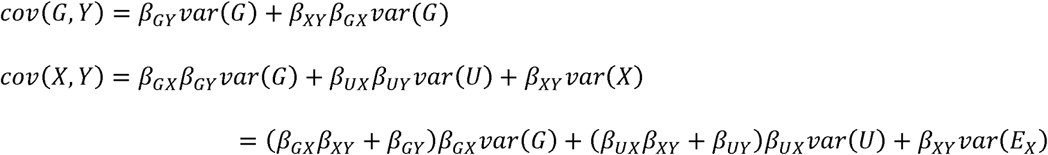

Substituting these covariances into Equation (5) gives, after some working out, Equation (3)

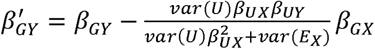

This derivation is similar to that of Aschard et al ^2^, except that we allow for the direct effect of *X* on *Y* in equation 2 and have focussed on the asymptotic estimate of the true 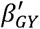.

As noted in Results, we may argue that 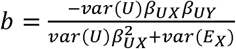 is approximately constant across SNPs and may be estimated by the linear regression of 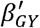 on /3_*GX*_ across many SNPs. In a finite sample this yields an estimate 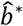 that is biased towards zero by sampling error in 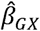. We suggest two approaches to adjust for this regression dilution bias. Firstly, following a common approach to the problem ^28^, we can obtain a bias-reduced estimate as 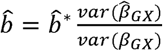. In the numerator 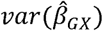 can be immediately estimated from the data, whereas estimation of *var*(/3_*GX*_) in the denominator is a well-studied problem in random effects meta-analysis ^35^. We find that the Hedges-Olkin estimator

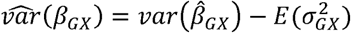

where 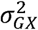 is the (estimated) sampling variance of 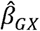, usually leads to acceptable estimates 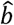 and given its ease of computation, we used this approach in our simulations.

However, as has been discussed in the context of Mendelian randomisation ^23^, this approach can have large variance, and can lead to implausible negative adjustments for regression dilution, as we found in our IPF data. Therefore we follow Bowden et al ^23^ in proposing simulation extrapolation ^24^ (SIMEX) for the analysis of real data sets. Briefly, this approach simulates new estimates 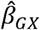 with increasing degrees of measurement error, by adding Gaussian noise with variance 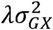 to the given 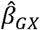, for various values of *λ*. The linear regression of Equation (3) is repeated for each simulated data set, and the estimator of its slope 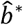 considered as a function of *λ*. Standard applications of SIMEX, including that of Bowden et al ^23^, fit a linear or quadratic model relating 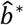 to *λ*, extrapolating to *λ* = −1 to obtain the de-biased estimate. For greater accuracy we developed a maximum likelihood estimator of b for simple linear regression models. Our approach yields confidence intervals for *b* so that 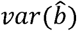 can be estimated. The details of our improved SIMEX approach are given in Supplementary Note 1.

The bias-corrected effect on prognosis is 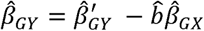, with variance

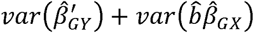

It is reasonable to assume that 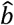 is approximately independent of 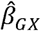 if a large number of independent SNPs enter the regression of Equation (3). Therefore

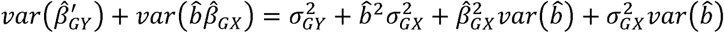

If 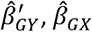 and 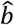 are maximum likelihood estimates, we may assume that they are approximately normally distributed about their true values with variance estimates available. As the product of two normal variates, 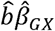 is not itself normal, but a bootstrap distribution for 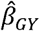 can be generated by simulating 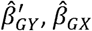 and 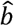 from their respective normal distributions, taking the estimated values as the mean. In the results presented we generally found that the bootstrap distribution was very close to normal and we therefore give *P*-values based on a normal approximation for 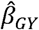. The exception was the analysis of rs35705950 in IPF, for which we simulated an empirical distribution for 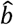 and then 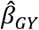 (Supplementary Note 1).

The derivation of Equation (3) assumes that *G, U, E*_*X*_ and *E*_*Y*_ are pairwise uncorrelated, which is unlikely to hold in general. Although by definition *U, EX* and *E*_*Y*_ are uncorrelated, *G* may be correlated with any of those variables through LD or gene-environment correlation. However, we might expect that across all SNPs in a GWAS, any systematic such correlation will be negligible. Equation (3) also assumes no statistical interaction between *G* and *U* in their effects on *X*, and between *G, U* and *X* in their effects on *Y*. Again, and in view of the low number of detectable interactions in GWAS compared to main effects ^36^, we may safely assume that any systematic interactions are negligible in comparison to the main effects.

The usual assumptions of linear regression apply to the estimation of *b*. The residuals, which are the mean-centred prognosis effects, should be uncorrelated. When marginal single-SNP effects are considered, as is usual in GWAS, correlation can arise through LD, and we therefore fit Equation (3) to a pruned set of approximately independent SNPs. Even with pruned SNPs, LD can lead to heteroscedasticity, since a SNP in a region of high LD is expected to have greater variance in its marginal effect on both incidence and prognosis ^37^. Furthermore, allele frequency has also been observed to relate to effect size variance ^38^, again creating potential heteroscedasticity. Residual heteroscedasticity does not affect the bias of 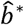 but its standard error is needed for our SIMEX adjustment, and so we calculate a heteroscedasticity robust estimate of that standard error (Supplementary Note 1).

Our most serious assumption is that residuals and predictor are uncorrelated in the regression: that is there is no correlation between a SNP’s effect on incidence *β*_*GX*_ and its direct effect on prognosis *β*_*GY*_. We discuss this assumption in the Results and Discussion and explore robustness to it in simulations.

Many GWAS will study prognosis among the cases of disease, rather than adjusting for an index trait as a covariate. The susceptibility GWAS will typically be performed using logistic regression, rather than linear regression as developed here. Index event bias has a non-linear form in logistic models but is approximately linear for the small effects typical of polygenic traits ^13^; furthermore, small effects on linear and logistic scales are approximately proportional ^39^. We therefore expect Equation (3) to hold approximately when *β*_*GX*_ and/or *β*_*GY*_ are log odds ratios. Having already assumed no interaction between *G* and *X* in their effects on *Y*, we further expect Equation (3) to hold when analysing only the cases of disease.

Finally, we have considered analyses with only genotype *G* and incidence *X* as predictors. In practice, further covariates will be included, such as principal components of ancestry. Analytic equivalents of Equation (3) are more complicated in this case, but one can often treat the conditional SNP effects as approximately equal to those on the residuals from a first-stage regression on the further covariates. With this justification we can apply our procedure to conditional SNP effects on incidence and prognosis.

### Simulations

SNPs were simulated independently under Hardy-Weinberg equilibrium with minor allele frequencies drawn uniformly from (0.01, 0.49). SNP effects, confounders and residual variation in incidence and prognosis were drawn independently from normal distributions. For heritability of 50% distributed among 10,000 SNPs with effects on prognosis, each SNP explains, on average, 0.005% of variation. As half of SNPs affecting prognosis also have effects on incidence, and the total non-genetic confounder variance is 40%, index event bias arises from confounders that together explain 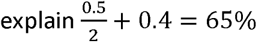 of variation in prognosis. Estimates of SNP effects on incidence 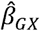 were obtained from linear regression of incidence on genotype, and unadjusted estimates of SNP effects on prognosis 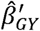 from linear regression of prognosis on genotype and incidence.

Incidence and prognosis traits were simulated from equations 1 and 2, with *β*_*GX*_ and *β*_*GY*_ now as the row vectors of effects for all SNPs, *G* as the column vector of genotypes and *U* consisting only of the non-genetic confounders.

We performed 1000 simulations and compared type-1 error at *P <* 0.05 for the unadjusted estimator 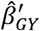 to our adjusted estimator 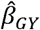, using the Hedges-Olkin estimator to correct for regression dilution. Type-1 error rates vary among SNPs since the index event bias is proportional to the effect on incidence and the rejection rate for a non-zero bias is greater for allele frequencies nearer 0.5. Firstly we estimated the mean type-1 error over all SNPs with no effect on prognosis. As this is dominated by the large number of SNPs without effects on incidence, and therefore no index event bias, we also estimated the mean type-1 error over SNPs with effects on incidence but not on prognosis. To assess lower error rates, we estimated the family-wise type-1 error over the same SNPs, as the proportion of simulations in which at least one SNP had *P* < 0.05 after Bonferroni adjustment for the number of such SNPs, that is *P* < 10^−5^. Finally we identified the individual SNP with the highest type-1 error for the unadjusted estimator and compared it to the type-1 error of our adjusted estimator for the same SNP.

Similarly, we estimated mean power over all SNPs with effects on prognosis, and all SNPs with effects on both incidence and prognosis. Equation (3) shows that index event bias may either increase or decrease power according to the particular values of its variables. Therefore, we identified the individual SNPs with the greatest increase and decrease in power between the unadjusted and proposed estimators.

We estimated bias and mean square error across SNPs in the same ways. Since we simulated genetic effects with mean zero, Equation (3) shows that the mean signed bias will be zero across SNPs, although individual SNPs will have non-zero bias. Therefore we estimated the mean absolute bias across SNPs.

We repeated the simulations with correlation between SNP effects on incidence and prognosis. For the 5000 SNPs with effects on both, we simulated their effects from a bivariate normal distribution with correlation 0.5, and then from a distribution with correlation 0.9. These led respectively to genome-wide genetic correlations between incidence and prognosis of 0.25 and 0.45. We repeated the simulations with the equivalent negative correlations.

We simulated a binary selection event by treating the incidence trait as a liability with a threshold for disease such that 20% of individuals were affected. We simulated 10,000 cases and 10,000 controls, and obtained estimated effects on incidence 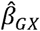 from logistic regression of disease on genotype. We then simulated a binary prognosis by thresholding the prognosis trait at its median, so that half the individuals had a good prognosis and half a poor prognosis. We obtained unadjusted estimated effects on prognosis 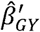 from logistic regression of prognosis on genotype among cases only.

For the binary selection event we also analysed the prognosis trait on its original quantitative scale using linear regression of prognosis on genotype among cases only, and compared results to the analysis of the combined case/control sample with statistical adjustment for case/control status, imputing a value of 0 for prognosis among controls. The latter approach may, in some situations, lead to reduced bias or increased power in comparison with case only analysis ^18^.

Genotype data and polygenic phenotypes were simulated using the --simulate and --score commands in PLINK 1.9 ^40^, with all other analyses performed in R 3.4.1.

### Idiopathic pulmonary fibrosis

612 cases and 3,366 controls previously genotyped in stage 1 of Allen et al ^26^ were used. Our secondary analysis is covered by the existing ethical approvals and informed consent reported for that study. Association with disease was adjusted for ten principal components of ancestry but not for age or sex, allowing inclusion of ten cases without data on age. Imputation was performed to the Haplotype Reference Consortium panel at the Michigan Imputation Server ^41^. We retained variants with imputation *R*^2^ of 0.5, minor allele frequency > 0.5%, Hardy-Weinberg equilibrium *P* > 10^−6^, and at least 5 events for subjects with allele dosage > 0.5. After harmonising the case/control and survival data we analysed 7,983,997 variants.

We created LD-pruned sets of SNPs using PLINK 1.9 ^40^ with *R*^2^ threshold of 0.1 within 250 SNP windows. To assess the effect of imputation quality on our procedure, we created separate pruned sets for SNPs with imputation *R*^2^ greater than 0.9, 0.98 and 0.99 in both incidence and survival GWAS. These sets contained 245,913, 154,095 and 140,092 SNPs respectively, and the regression of survival log hazard ratios on incidence log odds ratios gave coefficients of 0.048, −0.022 and −0.025 respectively. We were surprised to observe the change in sign of the coefficient as imputation *R*^2^ increased from 0.9 to 0.98, because the coefficient should be the same regardless of which SNPs are used for its estimation. Imputation introduces genotyping errors that do not follow a classical measurement error model because allele dosage is bounded in [0,2]. Furthermore, standard imputation methods do not take phenotype into account. As a result, effect sizes for imputed SNPs are biased in ways that have not been well characterised. While such biases must be small for well imputed SNPs, and have not created problems for standard GWAS analyses, the effect seems sufficient to bias our index event adjustment unless *R*^2^ >0.98 at least. Noting the compatibility between results for *R*^2^ >0.98 and 0.99, we used pruned SNPs meeting imputation *R*^2^ > 0.99 in both incidence and survival GWAS in all further analyses.

### Crohn’s disease

We downloaded summary statistics for incidence ^29^ (5,956 cases and 14,927 controls) and prognosis ^6^ (2,734 cases) from the internet and analysed 7,908,787 autosomal markers present in both datasets. Our re-analysis of these results is covered by the existing ethical approvals and informed consent reported for those studies. To estimate our regression adjustment we selected a set of 29,715 LD pruned SNPs, from a total of 1,370,154 SNPs with imputation *R*^2^ ≥ 0.99 in both the Crohn’s disease susceptibility GWAS and our IPF survival GWAS. LD was estimated using the genotypes of our IPF survival GWAS which has similar UK ancestry to the Crohn’s disease prognosis GWAS. The pruned set is smaller than those for IPF because the Crohn’s susceptibility GWAS was imputed to the 1000 Genomes reference, which yields fewer SNPs with high imputation *R*^2^ values than the Haplotype Reference Consortium reference.

### Code availability

An open source R package implementing the methods proposed in this report is available from https://github.com/DudbridgeLab/indexevent.

PLINK 1.9 is available from https://www.cog-genomics.org/plink2/

R 3.4.1 is available from https://cran.r-project.org/bin/windows/base/old/3.4.1/

### Data availability

The IPF data that support the findings of this study are available from the corresponding author upon reasonable request.

The Crohn’s disease susceptibility data that support the findings of this study are available from https://www.ibdgenetics.org/downloads.html

The Crohn’s disease prognosis data that support the findings of this study are available from ftp://ftp.sanger.ac.uk/pub/project/humgen/summary_statistics/human/2016-10-12/CD_prognosis_GWA_results.csv.zip.

All other data are available from the corresponding author upon reasonable request.

## Acknowledgements

Louise Wain holds a GSK / British Lung Foundation Chair in Respiratory Research. This article presents independent research funded partially by the UK National Institute for Health Research (NIHR). The views expressed are our own and not necessarily those of the NHS, the NIHR, or the UK Department of Health. Riyaz Patel is supported by a BHF Intermediate Clinical Research Fellowship (FS/14/76/30933).

## Author contributions

FD conceived the study, developed methods, performed analyses and wrote the manuscript. RJA performed analyses, interpreted results and edited the manuscript. NAS reviewed and edited the manuscript. AFS reviewed and edited the manuscript. JCL provided data, interpreted results, reviewed and edited the manuscript. RGJ provided data, reviewed and edited the manuscript. LVW provided data, interpreted results, reviewed and edited the manuscript. ADH conceived the study, interpreted results and reviewed the manuscript. RSP conceived the study, reviewed and edited the manuscript.

