## Supplemental Note and Tables for "Adjustment for index event bias in genome-wide association studies of subsequent events"

##### Dudbridge et al

### Supplementary Information

### Supplementary Note

#### 1. Simulation extrapolation

The idea behind simulation extrapolation (SIMEX) ^1,2^ is that if the bias in an estimator can be expressed as a function of measurement error, then an unbiased estimator can be obtained by setting the measurement error to zero. Suppose we have an estimate $\hat{b}^{*}$ for the regression slope of $\hat{\beta}_{GX}$, which itself is an unbiased estimate of $\beta_{GX}$ with measurement error being its sampling variance $\sigma_{GX}^{2}$. For a scalar $\lambda>0$ we can simulate new values $\hat{\beta}_{GX}^{(\lambda)}$ by adding Gaussian noise with variance ${\lambda\sigma}_{GX}^{2}$ to each observed value of $\hat{\beta}_{GX}$, and then obtain a (more biased) estimate $\hat{b}^{(\lambda)}$ for the regression slope of $\hat{\beta}_{GX}^{(\lambda)}$. If a functional form can be fitted to $\hat{b}^{(\lambda)}$ (note that $\hat{b}^{(0)}=\hat{b}^{*}$) then it can be extrapolated to provide an unbiased estimate as ${b=\hat{b}}^{(-1)}$. Usually, a limited range of values is considered for $\lambda$, and for each value $\hat{b}^{(\lambda)}$is taken as the mean over many simulations of the $\hat{\beta}_{GX}^{(\lambda)}$. In our analyses we ran 10,000 simulations for each $\lambda$ ranging from 0.25 to 5 in steps of 0.25.

A linear or quadratic model is typically fitted to $\hat{b}^{(\lambda)}$ ^3^. However we found that these models gave a poor fit to our idiopathic pulmonary fibrosis (IPF) data and therefore derived a maximum likelihood estimator of $b$ from data generated by SIMEX. From a standard result for the simple linear regression model ^4^

$$b=E\left( \hat{b}^{\left( \lambda\right)} \right)\frac{var\left( \hat{\beta}_{GX}^{\left( \lambda\right)} \right)}{var\left( \beta_{GX} \right)}=E\left( \hat{b}^{\left( \lambda\right)} \right)\left[ \frac{var\left( \hat{\beta}_{GX} \right)+\lambda\sigma_{GX}^{2}}{var\left( \beta_{GX} \right)} \right]$$

So

$$E\left( \hat{b}^{\left( \lambda\right)} \right)=b\left[ 1+(1+\lambda)\frac{\sigma_{GX}^{2}}{var\left( \beta_{GX} \right)} \right]^{-1}$$

(1)

This is written as an expectation because the actual $\hat{b}^{(\lambda)}$ obtained by SIMEX depend on the randomly simulated values of $\hat{\beta}_{GX}^{(\lambda)}$. To estimate the variance of $\hat{b}^{\left( \lambda\right)}$ we use the Huber-White sandwich estimator to allow for residual heteroscedasticity, as discussed in the main text. In general form this estimator is

$$\left( X^{T}X \right)^{-1}X^{T}\Sigma X\left( X^{T}X \right)^{-1}$$

where for now $X$ is the design matrix of the linear regression and $\Sigma$ is the diagonal matrix whose entries are the squared residuals of the regression. In the regression of $\hat{\beta}_{GY}$ on $\hat{\beta}_{GX}$ (Equation 3, main text) the design matrix consists of one column of 1’s (for the intercept) and a second column consisting of the $\hat{\beta}_{GX}$ for each SNP. The sandwich estimator is the 2×2 variance-covariance matrix of the estimated intercept and $\hat{b}^{\left( \lambda\right)}$, and we want the lower right entry. After some working out this is

$$\hat{var}\left( \hat{b}^{\left( \lambda\right)} \right)=\frac{\left( \sum\hat{\beta}_{GX}^{\left( \lambda\right)} \right)^{2}\sum U^{2}-2m\sum\hat{\beta}_{GX}^{\left( \lambda\right)}\sum\hat{\beta}_{GX}^{\left( \lambda\right)}U^{2}+m^{2}\sum{\hat{\beta}_{GX}^{\left( \lambda\right)}}^{2}U^{2}}{\left[ m\sum{\hat{\beta}_{GX}^{\left( \lambda\right)}}^{2}-\left( \sum\hat{\beta}_{GX}^{\left( \lambda\right)} \right)^{2} \right]^{2}}$$

(2)

where $m$ is the number of SNPs included in the regression of $\hat{\beta}_{GY}$ on $\hat{\beta}_{GX}^{(\lambda)}$, $U$ is the residual from this regression, and the sums are over the $m$ SNPs, with indices supressed for brevity.

Assume that each $\hat{b}^{(\lambda)}$ generated under SIMEX is normally distributed with the mean in Supplementary Equation 1 and variance in Supplementary Equation 2. If $\hat{b}^{(\lambda)}$ is taken as the mean over many simulations, then $\hat{var}\left( \hat{b}^{\left( \lambda\right)} \right)$ is correspondingly divided by the number of simulations. Then the log-likelihood for $b$ is

$$l\left( b;\frac{\sigma_{GX}^{2}}{var\left( \beta_{GX} \right)} \right)=\sum_{\lambda} \log{\phi(\hat{b}}^{\left( \lambda\right)};E\left( \hat{b}^{\left( \lambda\right)} \right),\hat{var}(\hat{b}^{\left( \lambda\right)}))$$

(3)

which we maximise over $b$ with $\frac{\sigma_{GX}^{2}}{var\left( \beta_{GX} \right)}$ as a nuisance parameter. This estimator gave a much better fit to our IPF data than the standard quadratic model, and a significantly different extrapolation (Supplementary Figure 1).

To obtain confidence intervals for $b$, we profile over $\frac{\sigma_{GX}^{2}}{var\left( \beta_{GX} \right)}$ as follows. The profile log-likelihood is as Supplementary Equation 3 with, for each value of $b$, $\frac{\sigma_{GX}^{2}}{var\left( \beta_{GX} \right)}$ replaced by the value that maximises the log-likelihood:

$$l_{b}\left( b \right)=l\left( b;\arg\max_{\theta} l(b;\theta) \right)$$

The $\left( 1-\alpha\right)\%$ confidence interval can be defined as the set of values $b$ that are not significantly different from the maximum likelihood estimate $\hat{b}_{\text{ML}}$ according to a likelihood ratio test of size $\alpha$. The confidence limits are then the solutions $b$ of

$$2\left( l_{b}\left( b \right)-l_{b}\left( \hat{b}_{\text{ML}} \right) \right)=\left[ \Phi^{-1}(\frac{\alpha}{2}) \right]^{2}$$

Under asymptotic normality of $\hat{b}_{\text{ML}}$, its variance can be inferred by dividing the difference between the confidence limits by ${2\Phi}^{-1}(1-\frac{\alpha}{2})$ which is approximately 3.92 in the usual case that $\alpha=0.05$.

If there is doubt over the normality of $\hat{b}_{\text{ML}}$, its empirical distribution can be estimated by simulating new $\hat{b}^{(\lambda)}$ for each $\lambda$ from the normal distribution with mean the actual $\hat{b}^{(\lambda)}$ and empirical variance from Supplementary Equation 2. From each set of simulated $\hat{b}^{(\lambda)}$, the maximum likelihood estimate $\hat{b}_{\text{ML}}$ is obtained to generate the empirical distribution of $\hat{b}_{\text{ML}}$. This may be combined with values of $\hat{\beta}_{GY}^{'}$ and $\hat{\beta}_{GX}$ simulated from asymptotic normal distributions to obtain the empirical distribution of $\hat{\beta}_{GY}=\hat{\beta}_{GY}^{'} -\hat{b}\hat{\beta}_{GX}$.

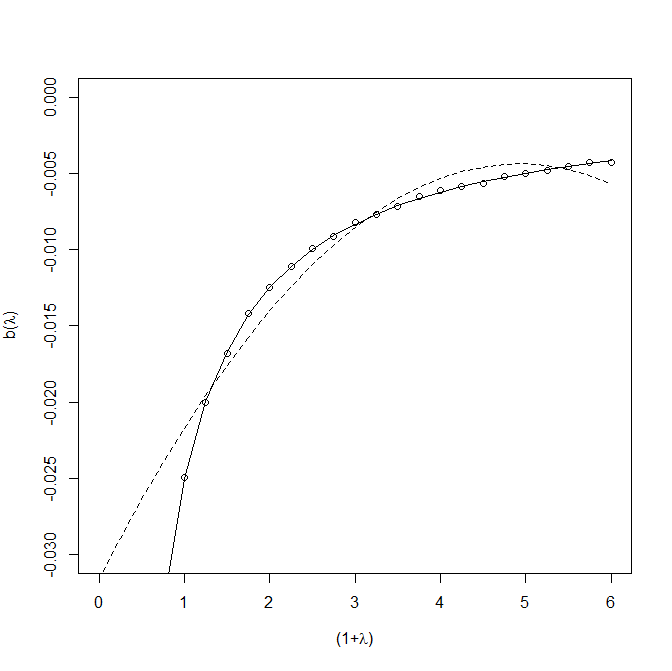

**Supplementary figure 1. SIMEX analysis of idiopathic pulmonary fibrosis data.**

Circles give the mean coefficient of the regression of $\hat{\beta}_{GY}$ on the simulated $\hat{\beta}_{GX}^{(\lambda)}$. The regression on the actual $\hat{\beta}_{GX}$ is shown at $(1+\lambda)=1$. Dotted line shows the quadratic fit obtained from standard software ^3^, which extrapolates to $\hat{b}^{(-1)}=-0.0316$. Solid line shows our maximum likelihood fit, which extrapolates to $\hat{b}^{(-1)}=-65.63$ (95% CI: -65.88, -5.68).

#### 2. Collider bias through selection on a subsequent trait

Under the directed acyclic graph of figure 2, we estimate the effect of $G$ on $X$ conditional on $Y$ as

$\left[ \begin{matrix} \beta_{GX}^{'} \\ \beta_{YX}^{'} \end{matrix} \right]=\left[ \begin{matrix} var\left( G \right) & cov\left( G,Y \right) \\ cov\left( G,Y \right) & var\left( Y \right) \end{matrix} \right]^{-1}\left[ \begin{matrix} cov\left( G,X \right) \\ cov\left( X,Y \right) \end{matrix} \right]$

$=\frac{1}{var\left( G \right)var\left( Y \right)-{cov(G,Y)}^{2}}\left[ \begin{matrix} var\left( Y \right) & -cov\left( G,Y \right) \\ -cov\left( G,Y \right) & var\left( G \right) \end{matrix} \right]\left[ \begin{matrix} cov\left( G,X \right) \\ cov\left( X,Y \right) \end{matrix} \right]$

Using the covariances given in the Methods, this yields

$\beta_{GX}^{'}=\frac{{var\left( U \right)(\beta}_{UY}+\beta_{UX}\beta_{XY})\beta_{UY}+var(E_{Y})}{{{var\left( U \right)(\beta}_{UY}+\beta_{UX}\beta_{XY})}^{2}+{var\left( E_{X} \right)\beta}_{XY}^{2}+var(E_{Y})}\beta_{GX}-\frac{{var\left( U \right)(\beta}_{UY}+\beta_{UX}\beta_{XY})\beta_{UX}+var(E_{X})\beta_{XY}}{var\left( U \right){{(\beta}_{UY}+\beta_{UX}\beta_{XY})}^{2}+var\left( E_{X} \right)\beta_{XY}^{2}+var(E_{Y})}\beta_{GY}$

While the bias is still linear in $\beta_{GY}$ with a slope that could be estimated from data, recovery of $\beta_{GX}$ requires knowledge of the confounder effects as well as the direct effect of $X$ on $Y$. Note however that if there is no such direct effect, then figure 1 and figure 2 are equivalent, and the above reduces to

$\beta_{GX}^{'}=\beta_{GX}-\frac{var\left( U \right)\beta_{UY}\beta_{UX}}{var\left( U \right){\beta_{UY}}^{2}+var(E_{Y})}\beta_{GY}$

analogous to Equation 3 in the main text.

### Supplementary tables

| Genetic correlation | 0 | | 0.25 | | 0.45 | | -0.25 | | -0.45 | |
| --- | --- | --- | --- | --- | --- | --- | --- | --- | --- | --- |
| Adjustment | No | Yes | No | Yes | No | Yes | No | Yes | No | Yes |
| All SNPs | 0.011 | 0.012 | 0.011 | 0.011 | 0.011 | 0.011 | 0.011 | 0.013 | 0.011 | 0.013 |
| All SNPs affecting incidence | 5.4e-3 | 2.4e-3 | 7.4e-3 | 4.7e-3 | 9.3e-3 | 7.5e-3 | 3.6e-3 | 4.8e-3 | 2.5e-3 | 7.4e-3 |

Supplementary Table 1. Absolute bias for quantitative incidence and prognosis with non-genetic confounding.

Estimates shown over 1000 simulations of100,000 independent SNPs. 5000 SNPs have effects on incidence only, 5000 on prognosis only and 5000 on both incidence and prognosis. Heritability of both incidence and prognosis is 50% with the genetic correlation shown over all SNPs. Common non-genetic factors explain 40% of variation in both incidence and prognosis.

| Genetic correlation | 0 | | 0.25 | | 0.45 | | -0.25 | | -0.45 | |
| --- | --- | --- | --- | --- | --- | --- | --- | --- | --- | --- |
| Adjustment | No | Yes | No | Yes | No | Yes | No | Yes | No | Yes |
| All SNPs | 2.4e-4 | 2.7e-4 | 2.4e-4 | 2.5e-4 | 2.4e-4 | 2.4e-4 | 2.3e-4 | 3.1e-4 | 2.4e-4 | 3.4e-4 |
| All SNPs affecting incidence | 2.5e-4 | 2.5e-4 | 2.8e-4 | 2.3e-4 | 3.2e-4 | 2.7e-4 | 2.4e-4 | 3.4e-4 | 2.4e-4 | 4.3e-4 |

Supplementary Table 2. Mean square error for quantitative incidence and prognosis with non-genetic confounding.

Parameters as in Supplementary Table 1.

| Genetic correlation | 0 | | 0.25 | | 0.45 | | -0.25 | | -0.45 | |
| --- | --- | --- | --- | --- | --- | --- | --- | --- | --- | --- |
| Adjustment | No | Yes | No | Yes | No | Yes | No | Yes | No | Yes |
| All SNPs | 0.011 | 0.011 | 0.011 | 0.011 | 0.011 | 0.011 | 0.011 | 0.013 | 0.011 | 0.011 |
| All SNPs affecting incidence | 2.4e-3 | 2.4e-3 | 3.4e-3 | 4.6e-3 | 4.7e-3 | 7.0e-3 | 3.4e-3 | 4.4e-3 | 4.7e-3 | 6.9e-3 |

Supplementary Table 3. Absolute bias for quantitative incidence and prognosis without non-genetic confounding.

Parameters are as in supplementary table 1 except that there are no common non-genetic factors of incidence and prognosis.

| Genetic correlation | 0 | | 0.25 | | 0.45 | | -0.25 | | -0.45 | |
| --- | --- | --- | --- | --- | --- | --- | --- | --- | --- | --- |
| Adjustment | No | Yes | No | Yes | No | Yes | No | Yes | No | Yes |
| All SNPs | 2.4e-4 | 2.4e-4 | 2.4e-4 | 2.5e-4 | 2.3e-4 | 2.5e-4 | 2.4e-4 | 2.4e-4 | 2.3e-4 | 2.4e-4 |
| All SNPs affecting incidence | 2.5e-4 | 2.5e-4 | 2.6e-4 | 2.8e-4 | 2.7e-4 | 3.4e-4 | 2.6e-4 | 2.7e-4 | 2.7e-4 | 3.3e-4 |

Supplementary Table 4. Mean square error for quantitative incidence and prognosis without non-genetic confounding.

Parameters are as in supplementary table 3.

| Genetic correlation | 0 | | 0.25 | | 0.45 | | -0.25 | | -0.45 | |
| --- | --- | --- | --- | --- | --- | --- | --- | --- | --- | --- |
| Adjustment | No | Yes | No | Yes | No | Yes | No | Yes | No | Yes |
| All SNPs | 0.037 | 0.038 | 0.041 | 0.041 | 0.046 | 0.046 | 0.035 | 0.036 | 0.034 | 0.036 |
| All SNPs affecting incidence | 0.021 | 0.018 | 0.021 | 0.020 | 0.021 | 0.021 | 0.020 | 0.018 | 0.019 | 0.017 |

Supplementary Table 5. Absolute bias for binary incidence and prognosis with non-genetic confounding.

Parameters as in Supplementary Table 1 with cases defined as subjects in the top 20^th^ percentile of the incidence trait, and poor prognosis as cases in the top 50^th^ percentile of the prognosis trait. Common non-genetic factors explain 40% of variation in both incidence and prognosis. Prognosis is analysed in cases only.

| Genetic correlation | 0 | | 0.25 | | 0.45 | | -0.25 | | -0.45 | |
| --- | --- | --- | --- | --- | --- | --- | --- | --- | --- | --- |
| Adjustment | No | Yes | No | Yes | No | Yes | No | Yes | No | Yes |
| All SNPs | 2.6e-3 | 2.7e-3 | 3.1e-3 | 3.2e-3 | 4.0e-3 | 4.0e-3 | 2.3e-3 | 2.5e-3 | 2.2e-3 | 2.4e-3 |
| All SNPs affecting incidence | 3.5e-3 | 3.6e-3 | 4.0e-3 | 4.1e-3 | 4.7e-3 | 4.8e-3 | 3.2e-3 | 3.2e-3 | 3.1e-3 | 2.9e-3 |

Supplementary Table 6. Mean square error for binary incidence and prognosis with non-genetic confounding.

Parameters as in Supplementary Table 5.

| Genetic correlation | 0 | | 0.25 | | 0.45 | | -0.25 | | -0.45 | |
| --- | --- | --- | --- | --- | --- | --- | --- | --- | --- | --- |
| Adjustment | No | Yes | No | Yes | No | Yes | No | Yes | No | Yes |
| All SNPs | 0.033 | 0.033 | 0.033 | 0.033 | 0.034 | 0.035 | 0.033 | 0.033 | 0.034 | 0.034 |
| All SNPs affecting incidence | 0.017 | 0.017 | 0.017 | 0.018 | 0.017 | 0.017 | 0.017 | 0.018 | 0.017 | 0.017 |

Supplementary Table 7. Absolute bias for binary incidence and prognosis without non-genetic confounding.

Parameters as in Supplementary Table 5 except that there are no common non-genetic factors of incidence and prognosis.

| Genetic correlation | 0 | | 0.25 | | 0.45 | | -0.25 | | -0.45 | |
| --- | --- | --- | --- | --- | --- | --- | --- | --- | --- | --- |
| Adjustment | No | Yes | No | Yes | No | Yes | No | Yes | No | Yes |
| All SNPs | 2.1e-3 | 2.1e-3 | 2.1e-3 | 2.1e-3 | 2.2e-3 | 2.3e-3 | 2.1e-3 | 2.1e-3 | 2.2e-3 | 2.2e-3 |
| All SNPs affecting incidence | 2.8e-3 | 2.8e-3 | 2.9e-3 | 2.8e-3 | 2.9e-3 | 2.8e-3 | 2.8e-3 | 2.8e-3 | 2.9e-3 | 2.8e-3 |

Supplementary Table 8. Mean square error for binary incidence and prognosis without non-genetic confounding.

Parameters as in Supplementary Table 7.

| Genetic correlation | | 0 | | | 0.25 | | 0.45 | | -0.25 | | -0.45 | |
| --- | --- | --- | --- | --- | --- | --- | --- | --- | --- | --- | --- | --- |
| Adjustment | | No | Yes | | No | Yes | No | Yes | No | Yes | No | Yes |
| All SNPs | Case only | 5.03 | 5.00 | | 5.07 | 5.03 | 5.11 | 5.08 | 5.01 | 5.02 | 5.00 | 5.05 |
|  | Case/control | 5.04 | 5.00 | | 5.07 | 5.02 | 5.11 | 5.09 | 5.02 | 5.02 | 5.01 | 5.05 |
| All SNPs affecting incidence | Case only | 5.66 | 5.09 | | 6.24 | 5.58 | 7.00 | 6.60 | 5.27 | 5.38 | 5.12 | 6.04 |
|  | Case/control | 5.64 | 5.02 | | 6.25 | 5.46 | 7.00 | 6.67 | 5.27 | 5.29 | 5.10 | 5.89 |
| SNP with highest error | Case only | 15.0 | 6.80 | | 23.5 | 12.9 | 37.4 | 30.4 | 9.90 | 10.2 | 8.00 | 15.9 |
|  | Case/control | 13.5 | 7.80 | | 22.3 | 11.6 | 30.4 | 26.9 | 10.0 | 9.10 | 7.80 | 17.7 |
| Family-wise error | Case only | 8.70 | | 5.30 | 13.0 | 9.70 | 21.2 | 17.8 | 5.00 | 5.60 | 5.20 | 10.9 |
|  | Case/control | 9.50 | | 5.10 | 12.7 | 7.70 | 22.6 | 19.2 | 6.50 | 5.90 | 5.90 | 11.3 |

Supplementary Table 9. Type-1 error for binary incidence and quantitative prognosis with non-genetic confounding.

Type-1 error shown as % at $P<0.05$. Parameters as in Supplementary Table 1 and cases defined as subjects in the top 20^th^ percentile of the incidence trait. Common non-genetic factors explain 40% of variation in both incidence and prognosis. Case only, prognosis analysed by linear regression among cases only. Case/control, prognosis set to zero for controls and analysed by linear regression in full sample with adjustment for case/control status.

| Genetic correlation | | 0 | | 0.25 | | 0.45 | | -0.25 | | -0.45 | |
| --- | --- | --- | --- | --- | --- | --- | --- | --- | --- | --- | --- |
| Adjustment | | No | Yes | No | Yes | No | Yes | No | Yes | No | Yes |
| All SNPs | Case only | 12.1 | 11.3 | 11.7 | 11.7 | 10.8 | 11.0 | 12.2 | 10.0 | 11.9 | 8.43 |
|  | Case/control | 12.1 | 11.4 | 11.7 | 11.7 | 10.8 | 11.0 | 12.1 | 10.1 | 11.9 | 8.57 |
| All SNPs affecting incidence | Case only | 12.3 | 11.3 | 10.8 | 11.0 | 8.04 | 8.56 | 13.0 | 9.62 | 12.8 | 7.02 |
|  | Case/control | 12.4 | 11.3 | 10.6 | 10.9 | 8.01 | 8.34 | 12.9 | 9.54 | 12.6 | 7.06 |
| SNP with greatest increase in power | Case only | 27.4 | 45.0 | 30.1 | 43.4 | 29.4 | 36.6 | 10.7 | 23.1 | 6.30 | 11.0 |
|  | Case/control | 30.4 | 47.8 | 26.1 | 39.1 | 39.6 | 45.3 | 19.8 | 34.6 | 8.50 | 12.3 |
| SNP with greatest decrease in power | Case only | 64.2 | 35.7 | 33.1 | 20.5 | 22.2 | 20.6 | 74.2 | 22.2 | 94.7 | 38.0 |
|  | Case/control | 52.0 | 25.6 | 32.9 | 22.0 | 11.3 | 9.90 | 65.1 | 20.7 | 72.9 | 17.5 |

Supplementary Table 10. Power for binary incidence and quantitative prognosis with non-genetic confounding.

Power shown as % at $P<0.05$. Parameters as in Supplementary Table 9.

| Genetic correlation | | 0 | | 0.25 | | 0.45 | | -0.25 | | -0.45 | |
| --- | --- | --- | --- | --- | --- | --- | --- | --- | --- | --- | --- |
| Adjustment | | No | Yes | No | Yes | No | Yes | No | Yes | No | Yes |
| All SNPs | Case only | 5.00 | 5.00 | 5.00 | 5.02 | 5.01 | 5.06 | 5.00 | 5.01 | 5.01 | 5.06 |
|  | Case/control | 5.00 | 5.00 | 5.00 | 5.02 | 5.01 | 5.06 | 5.00 | 5.01 | 5.01 | 5.06 |
| All SNPs affecting incidence | Case only | 5.01 | 5.03 | 5.05 | 5.35 | 5.19 | 6.13 | 5.06 | 5.33 | 5.20 | 6.10 |
|  | Case/control | 5.01 | 5.03 | 5.05 | 5.35 | 5.18 | 6.13 | 5.06 | 5.33 | 5.20 | 6.09 |
| SNP with highest error | Case only | 7.70 | 7.80 | 7.60 | 10.6 | 8.20 | 19.2 | 7.50 | 10.6 | 8.70 | 21.4 |
|  | Case/control | 7.70 | 7.90 | 8.00 | 10.3 | 8.30 | 19.2 | 7.50 | 10.5 | 8.60 | 21.4 |
| Family-wise error | Case only | 4.80 | 4.90 | 5.30 | 7.00 | 5.30 | 12.6 | 5.20 | 5.90 | 3.60 | 8.70 |
|  | Case/control | 5.00 | 4.80 | 4.80 | 7.40 | 5.50 | 12.2 | 5.20 | 5.80 | 3.80 | 9.10 |

Supplementary Table 11. Type-1 error for binary incidence and quantitative prognosis without non-genetic confounding.

Type-1 error shown as % at $P<0.05$. Parameters as in Supplementary Table 9 except that there are no common non-genetic factors of incidence and prognosis.

| Genetic correlation | | 0 | | 0.25 | | 0.45 | | -0.25 | | -0.45 | |
| --- | --- | --- | --- | --- | --- | --- | --- | --- | --- | --- | --- |
| Adjustment | | No | Yes | No | Yes | No | Yes | No | Yes | No | Yes |
| All SNPs | Case only | 11.0 | 10.9 | 10.8 | 10.4 | 10.3 | 9.16 | 10.8 | 10.4 | 10.3 | 9.20 |
|  | Case/control | 11.0 | 10.9 | 10.8 | 10.4 | 10.3 | 9.16 | 10.8 | 10.4 | 10.3 | 9.20 |
| All SNPs affecting incidence | Case only | 10.9 | 10.9 | 10.4 | 9.76 | 9.37 | 7.32 | 10.4 | 9.81 | 9.38 | 7.40 |
|  | Case/control | 10.9 | 10.9 | 10.4 | 9.76 | 9.37 | 7.33 | 10.4 | 9.81 | 9.37 | 7.41 |
| SNP with greatest increase in power | Case only | 13.3 | 15.1 | 13.8 | 20.2 | 7.40 | 10.2 | 9.50 | 15.6 | 4.80 | 7.50 |
|  | Case/control | 13.3 | 15.4 | 19.6 | 25.9 | 7.50 | 10.1 | 9.50 | 15.6 | 4.80 | 7.50 |
| SNP with greatest decrease in power | Case only | 31.7 | 30.2 | 50.3 | 37.2 | 68.8 | 40.0 | 55.4 | 42.8 | 65.3 | 38.2 |
|  | Case/control | 29.3 | 27.7 | 33.7 | 21.1 | 69.0 | 39.8 | 55.4 | 42.7 | 65.0 | 38.5 |

Supplementary Table 12. Power for binary incidence and quantitative prognosis without non-genetic confounding.

Power shown as % at $P<0.05$. Parameters as in Supplementary Table 11.

| Genetic correlation | 0 | | 0.25 | | 0.45 | | -0.25 | | -0.45 | |
| --- | --- | --- | --- | --- | --- | --- | --- | --- | --- | --- |
| Adjustment | No | Yes | No | Yes | No | Yes | No | Yes | No | Yes |
| All SNPs | 5.01 | 5.00 | 5.01 | 5.00 | 5.01 | 5.00 | 5.00 | 5.00 | 5.00 | 5.00 |
| All SNPs affecting incidence | 5.11 | 5.25 | 5.12 | 5.27 | 5.13 | 5.29 | 5.09 | 5.23 | 5.10 | 5.22 |
| SNP with highest error | 7.90 | 8.70 | 8.20 | 9.10 | 7.70 | 10.4 | 8.00 | 8.90 | 8.80 | 8.60 |
| Family-wise error | 7.50 | 7.70 | 7.90 | 7.90 | 7.10 | 6.90 | 7.10 | 7.20 | 8.70 | 8.10 |

Supplementary Table 13. Type-1 error for binary incidence and survival prognosis with non-genetic confounding.

Type-1 error shown as % at $P<0.05$. Parameters as in Supplementary Table 1 with cases defined as subjects in the top 20^th^ percentile of the incidence trait, and survival time simulated from the exponential model with the prognosis trait as the log hazard. Common non-genetic factors explain 40% of variation in both incidence and prognosis.

| Genetic correlation | 0 | | 0.25 | | 0.45 | | -0.25 | | -0.45 | |
| --- | --- | --- | --- | --- | --- | --- | --- | --- | --- | --- |
| Adjustment | No | Yes | No | Yes | No | Yes | No | Yes | No | Yes |
| All SNPs | 5.00 | 5.00 | 5.01 | 5.00 | 5.01 | 5.00 | 5.00 | 5.00 | 5.00 | 5.00 |
| All SNPs affecting incidence | 5.09 | 5.23 | 5.09 | 5.26 | 5.11 | 5.27 | 5.09 | 5.23 | 5.07 | 5.22 |
| SNP with highest error | 7.60 | 9.50 | 8.50 | 9.40 | 7.80 | 9.40 | 7.90 | 8.60 | 7.50 | 8.30 |
| Family-wise error | 7.80 | 7.70 | 8.40 | 7.40 | 9.50 | 7.50 | 8.50 | 7.50 | 7.70 | 8.10 |

Supplementary Table 14. Type-1 error for binary incidence and survival prognosis without non-genetic confounding.

Type-1 error shown as % at $P<0.05$. Parameters as in Supplementary Table 3 with cases defined as subjects in the top 20^th^ percentile of the incidence trait, and survival time simulated from the exponential model with the prognosis trait as the log hazard. There are no common non-genetic factors of incidence and prognosis.
